# Tuft cell-derived acetylcholine is an effector of type 2 immunity and directly targets helminth parasites in the gut lumen

**DOI:** 10.1101/2023.12.08.570757

**Authors:** Marième Ndjim, Fabien Herbert, Imène Gasmi, Charlène Joséphine, Sylvain Henri, Evgenia Turtoi, Steeve Thirard, Alicia Giordano, Claire Ciancia, Collette Britton, Eileen Devaney, Tom N. McNeilly, Sylvie Berrard, Andrei Turtoi, Rick M. Maizels, François Gerbe, Philippe Jay

## Abstract

Upon parasitic helminth infection, activated intestinal tuft cells secrete IL-25, which initiates a type 2 immune response during which *lamina propria* ILC2s produce IL-13. This causes epithelial remodelling, including tuft cell hyperplasia with an unknown function. We describe a novel cholinergic effector function of tuft cells, which we show are the only epithelial cells expressing Choline Acetyltransferase (ChAT). During parasite infections, mice with epithelial-specific deletion of ChAT have increased worm burden and faecal egg counts although they are able to mount a comparable type 2 immune response. Mechanistically, IL-13-amplified tuft cells release acetylcholine (ACh) into the gut lumen. We demonstrate a direct effect of ACh on worms, reducing their viability and fecundity via helminth muscarinic ACh receptors, with effects promoted by inhibition of acetylcholinesterase, an helminth-secreted enzyme. Thus, tuft cells are sentinels in naive mice, and their amplification upon helminth infections serves an additional type 2 immune response effector function.

## INTRODUCTION

Tuft cells are a cellular subset mostly found in digestive and respiratory epithelia, and play critical roles in mucosal host defence. In the intestinal epithelium, tuft cells are essentially known for their essential sentinel function during parasite infections. The presence of helminth or protozoa in the gut triggers tuft cell secretion of the alarmin cytokine IL-25 that initiates a type 2 immune response ^1–3^. Such a response is principally orchestrated by type 2 innate lymphoid cells (ILC2s) through the secretion of type 2 cytokines such as IL-4, IL-5 and IL-13. A critical aspect of IL-4 and IL-13 function is the profound remodelling they cause in the intestinal epithelium. This includes amplification of the mucus-producing goblet cells and the tuft cell lineages, as well as resistin-like beta (Retnlβ) ectopic expression by small intestinal goblet cells, in which it is usually absent ^4^. While Retnlβ directly interferes with worm physiology ^4–6^, increased mucus production and smooth muscle hypercontractility, also known as the “weep and sweep” response, facilitates worm expulsion ^7^. In contrast, the physiological role of the dramatic increase in tuft cell numbers during type 2 immune responses is not yet understood. Increased IL-25 production following tuft cell lineage amplification is thought to lead to a more efficient type 2 immune response ^1,3^, but alternative tuft cell functions also need to be considered. In particular, we note firstly that this amplification occurs downstream of the action of type 2 cytokines on epithelial cells, and secondly that worm expulsion is significantly more delayed by the absence of tuft cells ^1^ as compared to IL-25 deficiency alone. These points indicate that in addition to their alarm function ^8^, tuft cells are required, not as initiators, but rather as an integral effector component of the type 2 immune response.

The acetylcholine (ACh) neurotransmitter, biosynthesized by the choline acetyl transferase (ChAT) enzyme, regulates a variety of neuronal and non-neuronal physiological functions ^9^. Studies with ChAT-reporter mice revealed the presence of cholinergic epithelial cells in various tissues including the trachea ^10^, urethra ^11^, gastro-intestinal and biliary tracts ^12^, and thymus ^13^. These cells were identified as tuft (also called brush or solitary chemosensory) cells in the airway, where their actual ACh release was demonstrated ^14^. In the nasal cavity, cholinergic tuft cells regulate breathing and inflammation in the presence of irritants ^15^; and in the trachea, they control breathing reflexes ^16^ and muco-ciliary clearance in response to bacterial quorum sensing molecules ^17^. In addition, urethral tuft cells control micturition reflexes through cholinergic signalling to viscerosensory neurons in response to exogenous bitter compounds ^11^. In contrast, the function of intestinal ChAT-expressing cells remains unknown.

Interestingly, some commonly used drugs against intestinal helminths, such as levamisole and pyrantel are cholinergic agonists. They target the worm acetylcholine receptors (AChRs), causing spastic paralysis which facilitates worm expulsion ^18^, and suggests potential direct effects of host ACh on parasites. In addition, in the gut mucosa, available acetylcholine for signalling results from the balance between its synthesis by the ChAT enzyme and its breakdown catalysed by acetylcholinesterase (AChE) enzymes. It is striking that, in addition to the neuromuscular AChE enzymes found in many organisms, some parasitic nematodes that colonize mucosal surfaces, encode additional AChE isoforms, which are produced in specific secretory glands and secreted into the worm environment ^19^, most likely to avoid detrimental exposure to host acetylcholine.

Here, we investigated the specific role of tuft cell-derived ACh in the context of type 2 immune responses. We found that tuft cells are the only intestinal epithelial cells expressing the key choline acetyl transferase (ChAT) enzyme for ACh biosynthesis, and confirmed their production of ACh. Worm clearance was delayed in mice with *ChAT*-deficient intestinal epithelial cells, in spite of the establishment of a strong type 2 immune response. Mechanistically, we demonstrated luminal production of ACh by tuft cells, and a direct effect of ACh on worms, resulting in reduced fitness and fecundity *via* a muscarinic acetylcholine receptor-dependent pathway.

## RESULTS

### The ACh biosynthesis enzyme ChAT is specifically expressed by tuft cells in the intestinal epithelium

We directly assessed the specific potential of small intestinal tuft cells to biosynthesize the acetylcholine neurotransmitter by analysing expression of the choline acetyl transferase (ChAT) enzyme that catalyses biosynthesis of acetylcholine in cultured intestinal organoids. Because tuft cells are poorly represented in normal mouse small intestinal organoids, we exposed the cultures to IL-13 to amplify the tuft cell lineage prior to analysis of tuft cell gene expression. *Chat* mRNA expression was found in wildtype organoids, which contain all epithelial cell subsets, including tuft cells. In contrast, *Chat* expression could not be detected in organoids derived from mice lacking the Pou2f3 transcription factor, which is required for tuft cell differentiation ^20,1^, strongly suggesting tuft cell-specific expression of ChAT (Figure 1A). Expression of *Pou2f3* mRNA was used as a control, and was detected in wildtype organoids but not in *Pou2f3*-deficient organoids (Figure 1B). To confirm this finding in the context of an entire organism, small intestinal EpCam+;Siglec-F+ tuft cells were enriched by FACS. Efficiency of the cell sorting procedure was assessed by analysing the expression of the tuft cell marker *Trpm5* mRNA in the tuft and non-tuft cell fractions. The *Trpm5* mRNA was detected in the tuft cell fraction whereas it was below the threshold of detection in the fraction containing non-tuft cells (Figure 1C). Similarly, *Chat* expression was detected in the tuft cell fraction but not in the fraction containing all non-tuft cell epithelial subsets (EpCam+;Siglec-F-) of the small intestinal epithelium, indicating specific expression of *Chat* in tuft cells among intestinal epithelial cells in vivo (Figure 1D). Finally, to assess whether *Chat* mRNA expression is a common property of small intestinal tuft cells or is limited to only a subset of these cells, in the context of an intact tissue, we coupled anti-Dclk1 immunofluorescence detection of the entire tuft cell population to *in situ* hybridization to visualize *Chat*-expressing cells in naive C57BL/6 mice. This revealed specificity of the *Chat* probe signal in Dclk1-expressing cells. Furthermore, almost all Dclk1-expressing tuft cells also displayed *Chat* mRNA signals (264/278 examined cells) in the proximal and distal small intestines as well as in the colon (Figure 1E and Supplementary data Figure 1).

**Figure 1:**
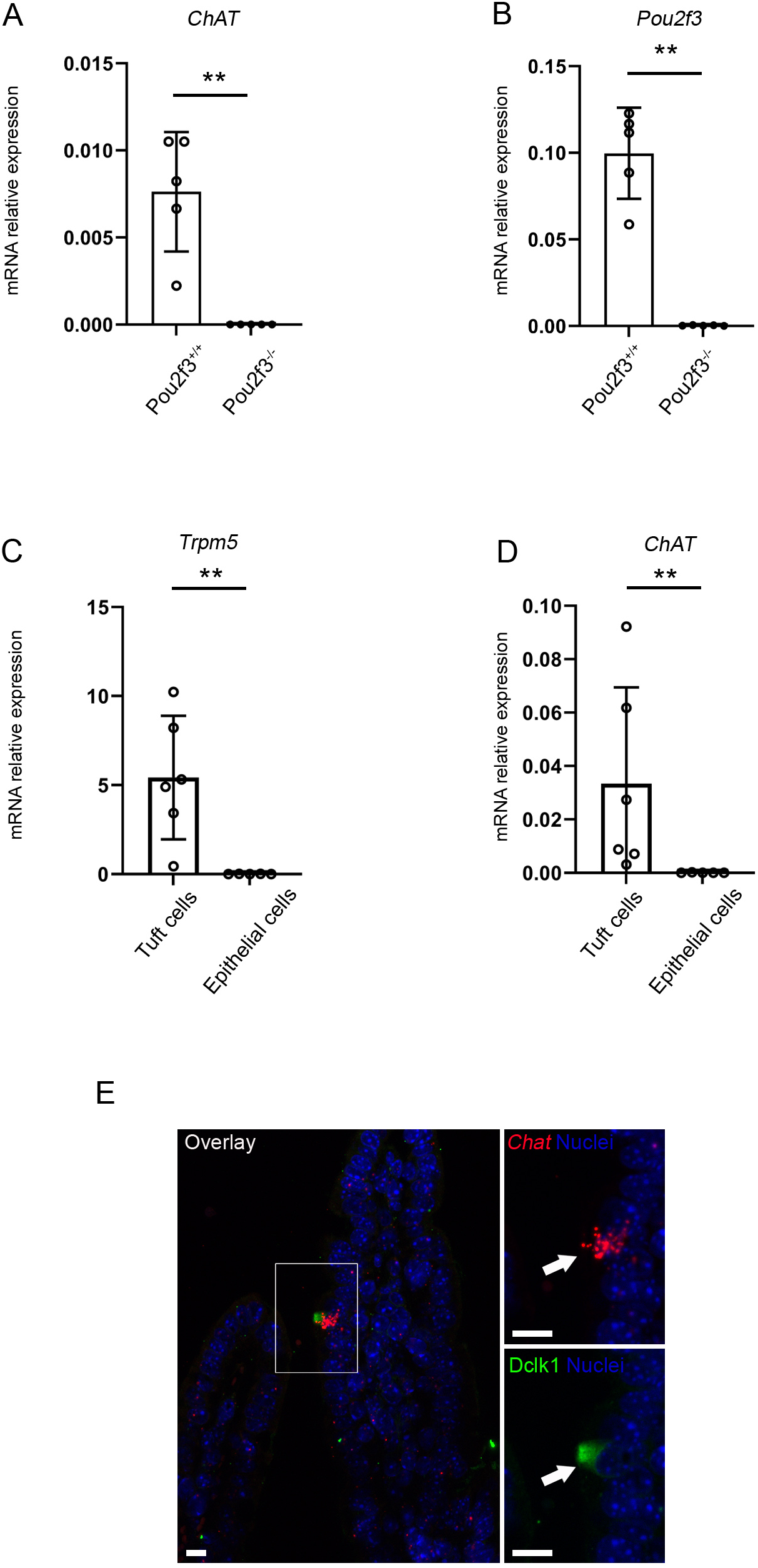
ACh production is restricted to tuft cells in the intestinal epithelium. (A-B) Expression of *ChAT* (A) and *Pou2f3* (B) transcripts in control or *Pou2f3*-deficient organoids treated with IL-13 for 72 hours. Data represent means ± SD of the biological replicates (n=5 for each genotype, with ** = p < 0.01 after a Mann Whitney test). (C-D) Expression of *Trpm5* (C) and *ChAT* (D) transcripts in FACS-sorted tuft cells as compared to all other epithelial cells, from naïve C57BL/6J mice. Bars represent means ± SD of the biological replicates (n=6 for tuft cell fractions, n=5 for all other epithelial cell fractions, with ** = p < 0.01 after a Mann Whitney test. (E) Representative *in situ* hybridisation for *Chat* mRNA (red)-coupled with immunofluorescence detection of Dclk1 (green). Scale bar = 10 µm for all panels. Arrows point at tuft cells positive for the *Chat* signal.

### Maintenance of ChAT expression during type 2 immune responses

We then asked whether ChAT expression is also a common feature of the amplified tuft cell population in the context of an helminth parasite infection ^1,3^. Wild type mice were thus infected by gavage with *Heligmosomoides polygyrus* infective L3 larvae which, after reaching the small intestine, penetrate the submucosa. There, they undergo two developmental moults before emerging again around day 10 post infection, as adult worms, into the gut lumen where they mate and produce eggs which are passed out in the faeces. The presence of *H. polygyrus* worms in the gut lumen is associated with a strongly polarised type 2 immune response ^21^, which includes dramatic amplification of tuft cells. *Chat* mRNA was also found in the majority of tuft cells from *H. polygyrus*-infected mice (Figure 2A). Furthermore, a quantitative comparison revealed similar proportions of *Chat*^+^ Dclk1-expressing tuft cells in naive and infected mice (Figure 2B). Thus, *Chat* mRNA expression is present specifically in tuft cells in the mouse small intestinal epithelium and can be detected in almost all tuft cells regardless of their infection status.

**Figure 2:**
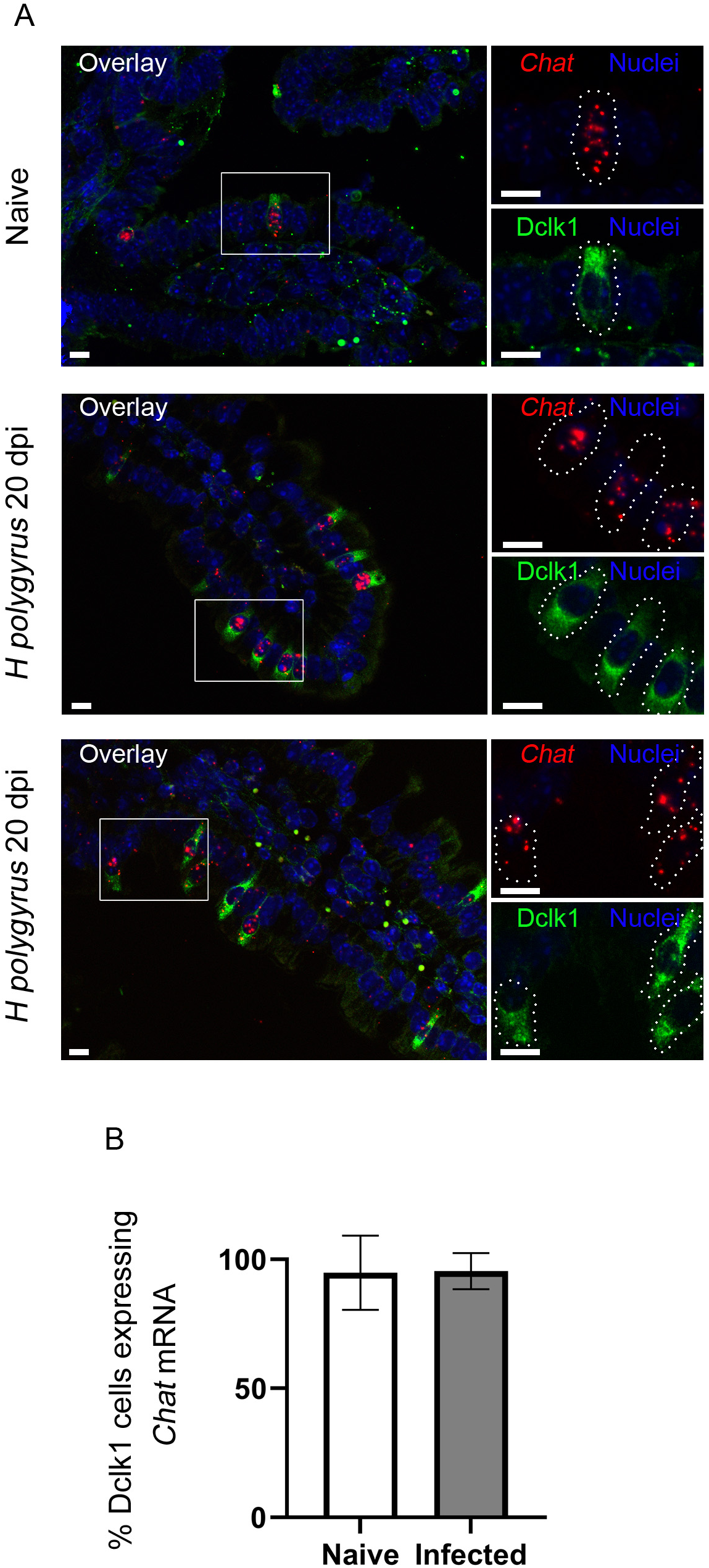
*ChAT* expression is restricted to tuft cells during *H. polygyrus* infection. (A) Representative *in situ* hybridisation for *Chat* mRNA (red)-coupled with immunofluorescence detection of Dclk1 (green). Top panel: representative picture of a small intestinal section from a naive mouse; middle and bottom panel, representative pictures from twi replicate infected mice at day 20 post-infection. Dotted lines highlight tuft cells co-expressing Dclk1 and *ChAT* mRNA. Scale bar = 10 µm for all panels. (B) Percentage of Dclk1-expressing cells co-expressing *Chat* mRNA in naive (white bar, n=3) and *H. polygyrus* infected (grey bar, n=2) conditions.

### Impaired host defence against helminth parasites in ChAT-deficient mice

To investigate the function of tuft cell-produced ACh in the context of an *in vivo* type 2 immune response, we generated an inducible deletion of the *Chat* gene specifically in the intestinal epithelium by crossing *Chat^LoxP/^*^LoxP^ ^22^ and *Villin-Cre^ERT2^* mice, to express Cre recombinase in a tamoxifen-inducible manner ^23^. Since the ChAT gene is likely expressed in intestinal non epithelial cells, we checked the correct recombination of the *LoxP* sites in organoids, constituted only of epithelial cells, derived from *Chat^LoxP/LoxP^;Villin-Cre^ERT2^* mice treated with tamoxifen (Figure 3A). We also confirmed the expression of the Cre^ERT2^ fusion protein in differentiated tuft cells (Supplementary data Figure 2). In addition, because the *Villin* gene promoter is active in all intestinal epithelial cells, including stem cells ^23^, Cre^ERT2^ activation by tamoxifen causes permanent gene deletion in compound *Chat^LoxP/LoxP^;Villin-Cre^ERT2^* mice. No gross alteration of the epithelial layer was perceptible as identical representations of epithelial cells from the enteroendocrine, Paneth and goblet lineages were found in wild type mice and mice with ChAT-deficient epithelial cells (Supplementary data Figure 3). To assess the consequences of the ChAT deficiency during an infection with parasitic helminths, we first induced *chat* gene deletion by tamoxifen treatment during five consecutive days. After 5 days of rest without tamoxifen, *Chat^LoxP/^*^LoxP^ and *Chat^LoxP/LoxP^;Villin-Cre^ERT2^* mice were infected by gavage with *H. polygyrus* L3, and infection parameters were analysed at 20 and 40 days post-infection when adult worms are present in the gut lumen (Figure 3B). Although *Chat*-deficient mice had no obvious phenotype, autopsy revealed increased numbers of adult worms in these mice as compared to *Chat^LoxP/LoxP^* control mice (Figure 3C). Greater numbers of live worms were mirrored by the numbers of eggs in the faeces of infected mice, a dynamic readout of the type 2 immune response efficiency and worm persistence and fitness. Increased numbers of eggs were found in *Chat*-deficient mice between 10 days post-infection, when adult worms start to be present in the gut lumen, and 40 days post-infection, as compared to *Chat^LoxP/LoxP^* littermates (Figure 3D). Increased numbers of adult worms and faecal eggs in *Chat*-deficient mice thus revealed an essential role of tuft cell-derived acetylcholine for an efficient response against helminth parasites.

**Figure 3:**
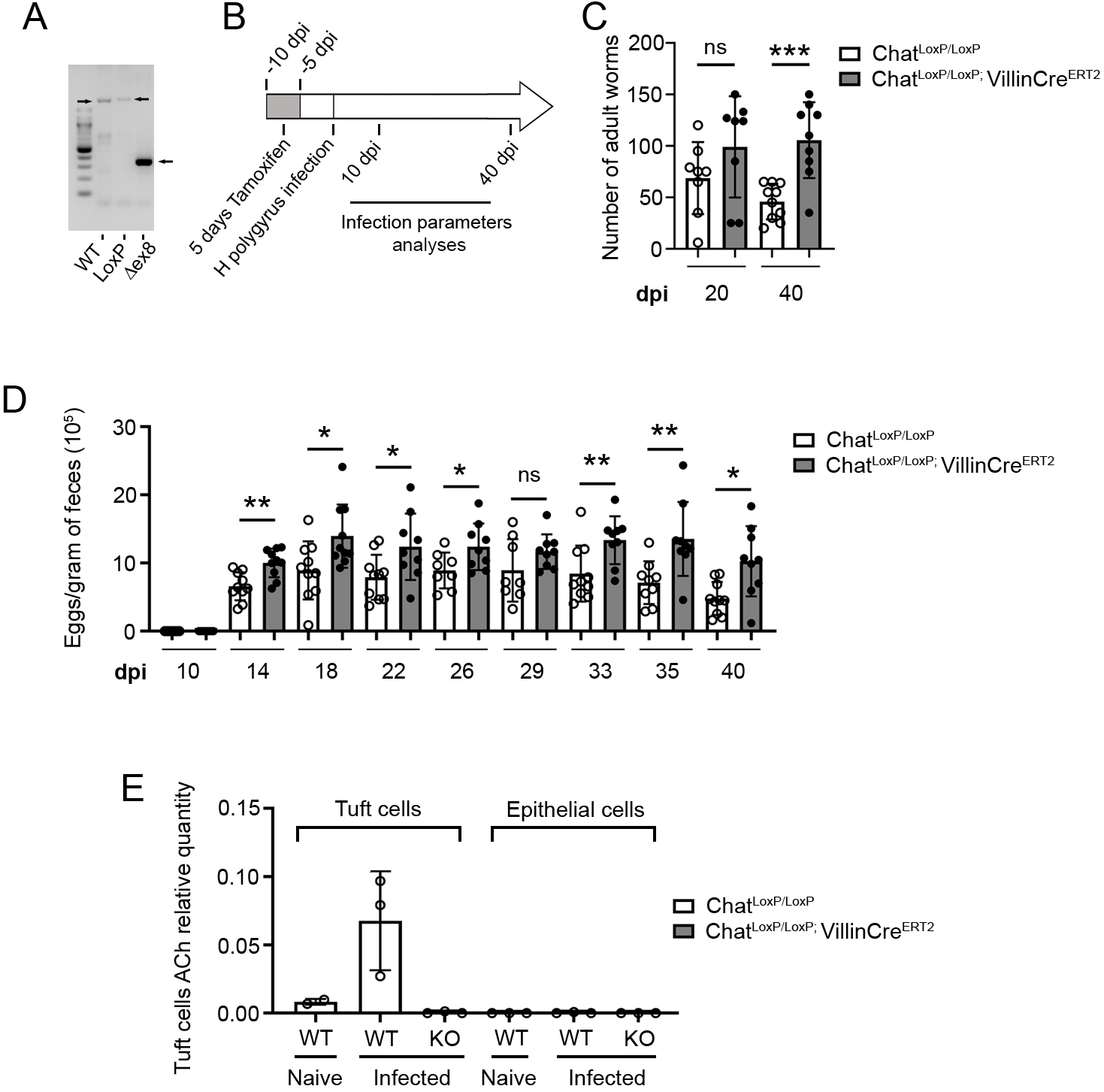
Tuft cell ChAT deficiency delays worm expulsion. (A) Efficient *Chat* locus recombination shown by PCR amplification of genomic DNA obtained from tamoxifen-treated organoids derived from *Chat^LoxP/LoxP^* or *Chat^LoxP/LoxP^;Villin-Cre^ERT2^*mice, using primers flanking exon 8 of the genetically engineered *Chat* locus. Arrows point bands at the expected sizes. (B) Scheme depicting the experimental design of *H. polygyrus* infections and mouse analysis time points. Control mice (*Chat^LoxP/LoxP^*) or mice with an intestinal epithelium-targeted *Chat* gene deletion (*Chat^LoxP/LoxP^;Villin-Cre^ERT2^*) were first treated with 1 mg of tamoxifen between −10 and −5 dpi, and then orally infected with *H. polygyrus* L3 larvae at day 0. Infected animals were monitored and/or sacrificed from 10 to 40 days post-infection. (C) Kinetics of infection of *Chat^LoxP/LoxP^* (white bars and circles) or *Chat^LoxP/LoxP^;Villin-Cre^ERT2^* mice (dark bars and circles) with 200 *H. polygyrus* L3, monitored for numbers of adult worms found in their small intestines. Data represent means ± SD of the different biological replicates (n ranging from 5 to 10 mice per time point, *** = p < 0.001, after a Mann Whitney test). (D) Kinetics of infection of *Chat^LoxP/LoxP^* (white bars and circles) or *Chat^LoxP/LoxP^;VillinCre^ERT2^* mice (dark bars and circles) with 200 *H. polygyrus* L3, monitored for numbers of eggs per gram of faeces. Data represent means ± SD of the different biological replicates (n ranging from 5 to 10 mice per time point, * = p < 0.05, ** = p < 0.01, after a Mann Whitney test). (E) Mass spectrometry-based acetylcholine relative quantification in FACS-sorted tuft cells as compared to all other epithelial cells isolated from either control *Chat^LoxP/LoxP^* (WT) or *Chat^LoxP/LoxP^;VillinCre^ERT2^* (KO) mice, in naïve and *H. polygyrus*-infected contexts. Bars represent means ± SD of different biological samples (n=2 to 3 cell fractions per group).

### Increased luminal ACh released by tuft cells during *H. polygyrus* infection

We then assessed the tuft cell production of ACh by mass spectrometry using lysates from FACS-sorted tuft and non-tuft cell fractions. We found a striking elevation of ACh In the EpCam+;Siglec-F+ tuft cell fractions of mice infected with *H. polygyrus*, as compared to the tuft cell fractions from naïve mice, although the limited number of replicates for the naïve condition precluded descriptive statistical analysis (Figure 3E). As this elevation occurred in cellular populations enriched in tuft cells, it likely reflects increased ACh synthesis in tuft cells rather than a consequence of increased tuft cell numbers caused by the epithelial remodelling occurring in the context of type 2 immune responses. As expected, ACh was not detected in the tuft cell fractions of ACh-deficient *Chat^LoxP/LoxP^;Villin-Cre^ERT2^* mice, nor in the EpCam+;Siglec-F-non-tuft cell epithelial fractions from either *Chat^LoxP/LoxP^* control mice or *Chat^LoxP/LoxP^;Villin-Cre^ERT2^* ACh-deficient mice (Figure 3E).

### ChAT-deficient mice are able to mount a type 2 immune response

To understand the mechanisms leading to decreased parasite clearance in *ChAT^LoxP/LoxP^;Villin-Cre^ERT2^* mice, we assessed critical parameters of the type 2 immune response and subsequent epithelial remodelling in naive and *H. polygyrus*-infected mice. All the studied parameters of the type 2 immune response, including numbers of Gata3+ ILC2s/Th2 cells, epithelial tuft and goblet cells and expression of the Retnlβ peptide by small intestinal goblet cells indicated the presence of a strong type 2 immune response in both *Chat^LoxP/LoxP^*control mice and *Chat^LoxP/LoxP^;Villin-Cre^ERT2^* ACh-deficient mice, with some of these parameters reaching even higher levels in ACh-deficient infected mice, possibly due to the presence of higher numbers of worms (Figure 4A, 4B, 4C and Supplementary data Figure 4). The slightly decreased numbers of Gata3+ ILC2s/Th2 and tuft cells observed in naive ChAT- deficient animals likely reflect a role of tuft cell ACh at the steady state (Figure 4A). Together, these data indicate that, although *Chat^LoxP/LoxP^;Villin-Cre^ERT2^* ACh-deficient mice have slightly lower basal levels of some type 2 immune parameters, they are able to mount a strong type 2 immune response in the absence of tuft cell-derived acetylcholine. Therefore, higher worm persistence and egg production in *ChAT*-deficient mice is not due to a compromised type 2 immune response, indicating a yet unappreciated role of ACh in promoting worm expulsion.

**Figure 4:**
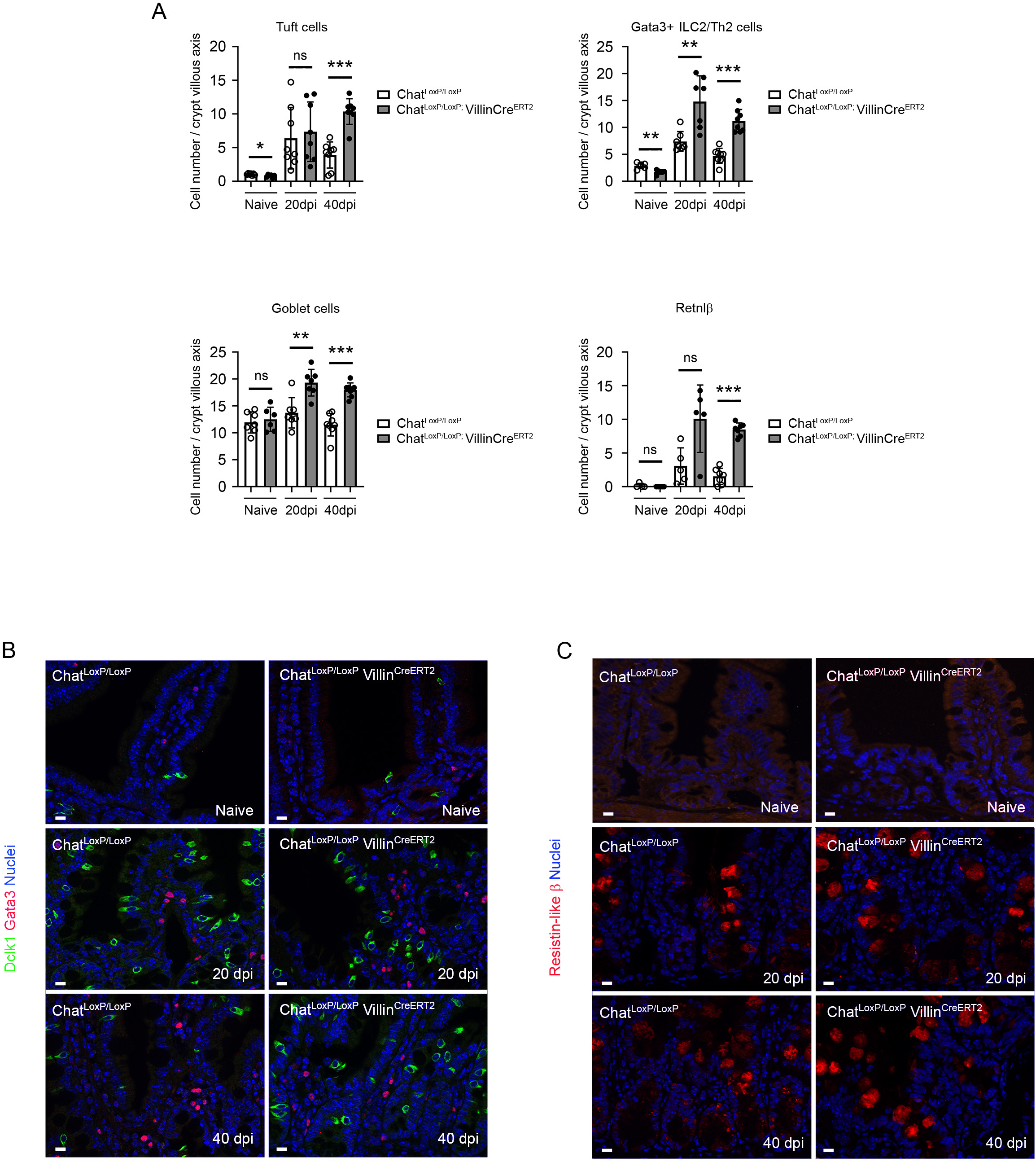
ChAT deficiency does not impair establishment of a type-2 immune response. (A) Characterisation of the intestinal mucosa of *Chat^LoxP/LoxP^* (white bars and circles) and *Chat^LoxP/LoxP^;VillinCre^ERT2^* mice (dark bars and circles) following *H. polygyrus* infection, at the indicated time points. Each cell type was quantified by IHC, using markers for tuft cells (Dclk1), ILC2s and Th2 immune cells (Gata3), mucus-secreting goblet cells (alcian blue and resistin-like β). Data represent means ± SD of n = 50 villus – crypt axes of each biological replicate (n ranging from 6 to 7; *= p < 0.05, ** = p < 0.01, *** = p < 0.001 after a Mann Whitney test). (B) Representative co-immunostainings of Dclk1 (green), Gata3 (red) and nuclei (blue) in naïve or infected *Chat^LoxP/LoxP^* and *Chat^LoxP/LoxP^;VillinCre^ERT2^* mice, 20 and 40 days post infection. (C) Representative co-immunostainings of Resistin-like β (red) and nuclei (blue) in naïve or infected *Chat^LoxP/LoxP^* and *Chat^LoxP/LoxP^;VillinCre^ERT2^* mice, 20 and 40 days post infection. Scale bars = 10 µm for B and C.

### Increased luminal ACh concentrations in *H. polygyrus*-infected mice

To investigate the hypothesis of a direct effect of tuft cell-derived ACh on worms present in the gut lumen, we assessed ACh production by intestinal epithelial cells. Comparison of ACh concentrations in the culture medium of intestinal organoids treated or not with the key type 2 cytokine IL-13 did not reveal significant differences (Supplementary data Figure 5), suggesting predominantly luminal production of ACh in IL-13-treated organoids (of note, intestinal organoids have an inverted polarity, meaning that the cellular apical side faces the organoid lumen).

We thus directly assessed luminal ACh relative concentrations in naive and *H. polygyrus*-infected mice. For this, an intestinal loop was surgically ligatured and filled with wash buffer, which was recovered after 30 minutes of incubation, and processed for detection of ACh by mass spectrometry. This revealed elevated ACh concentration in the gut lumen of wildtype infected mice, as compared to naive animals (Figure 5A), which is likely due to contribution of both higher tuft cell numbers and increased ACh production as suggested previously (Figure 3E). ACh concentrations were variable in infected mice, probably reflecting infection efficiencies as well as the specific location of worms as regards the favourable intestinal loop for surgery. Thus, increased concentrations of ACh are present in the gut lumen of infected mice and may directly impact worm physiology.

**Figure 5:**
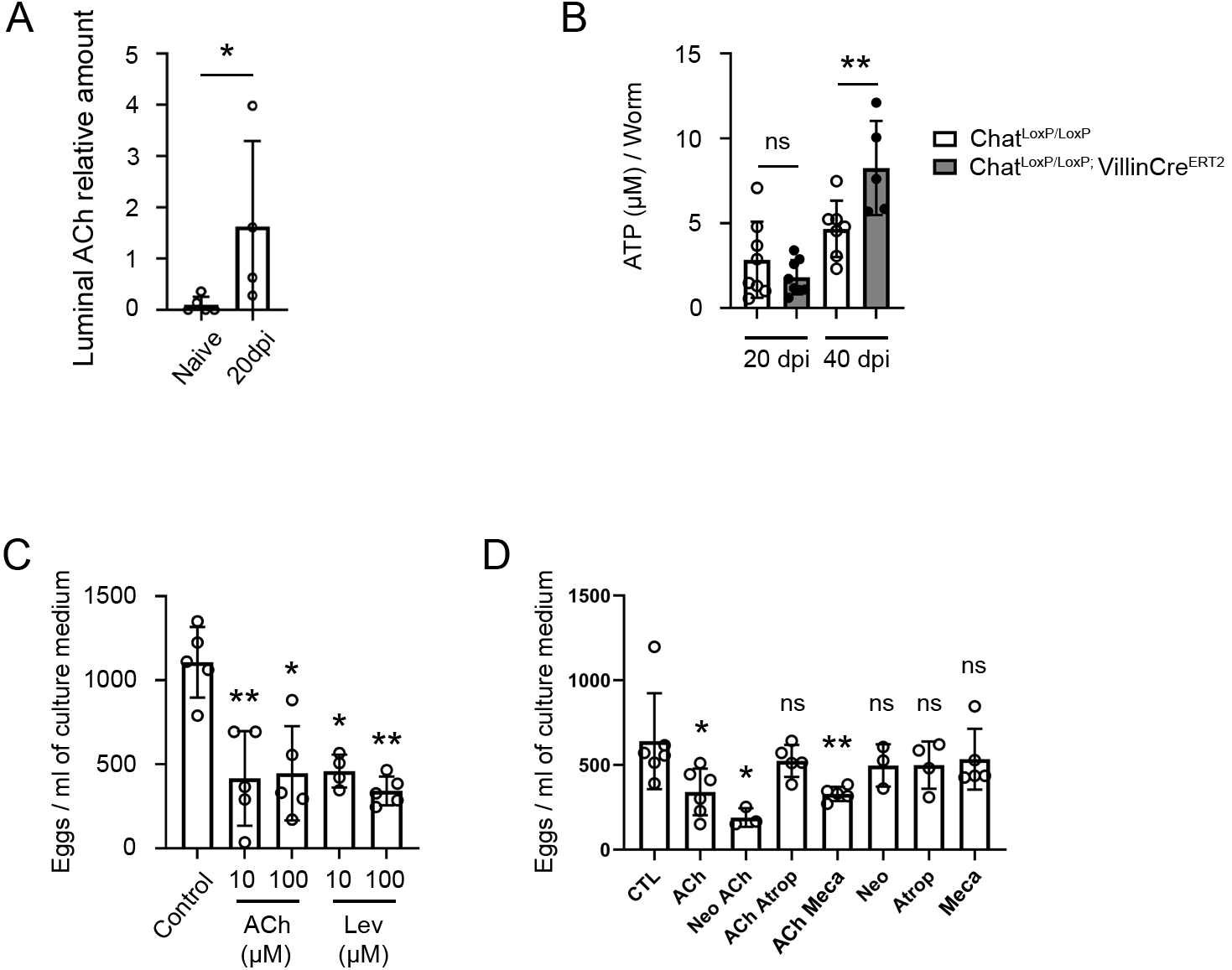
Tuft cell-derived ACh is released into the host lumen, and directly impairs worm physiology. (A) Quantification of luminal ACh, as described in the methods section, in naïve and *H. polygyrus*-infected C57BL/6J mice, 20 days post infection. Data represent means ± SD of the different biological replicates (n ranging from 4 to 5 mice, * = p < 0.05, after a Mann Whitney test). (B) ATP levels (µM) in worms recovered from either *Chat^LoxP/LoxP^* (white bars and circles) or *Chat^LoxP/LoxP^;Villin-Cre^ERT2^* mice (dark bars and circles) infected with *H. polygyrus* at different time points. Data represent means ± SD of the different biological replicates (n ranging from 4 to 8 groups of 25 worms per time point, ** = p < 0.01, after a Mann Whitney test). (C) Numbers of eggs released in culture medium during a 24h time frame, after treatment of adult *H. polygyrus* worms with acetylcholine (ACh) or levamisole (Lev) at 10 µM or 100 µM, as compared to untreated worms. Data represent means ± SD of the different biological replicates (n=5 groups of 25 female worms per condition, * = p < 0.05, ** = p < 0.01, after a Mann Whitney test). (D) Number of eggs released in culture medium during a 24h time frame after treatment of *H. polygyrus* adult worms with 10 µM acetylcholine (ACh), combined or not with 5 µM Neostigmine (Neo), 5 µM Atropine (Atrop), or 5 µM Mecamylamine (Meca), as compared to untreated worms. Data represent means ± SD of the different biological replicates (n ranging from 3 to 6 groups of 25 female worms per condition, * = p < 0.05, ** = p < 0.01 after a Mann Whitney test).

### Increased worm fitness and reproductive potential in ACh-deficient tuft cells

We next assessed the consequences of *H. polygyrus* exposure to ACh by two complementary approaches. Noteworthily, intestinal goblet cells produce another molecule, Retnlβ, specifically in the context of type 2 immune responses, and this directly interferes with *H. polygyrus* physiology ^4^. Subsequently, it was found that worms exposed to Retnlβ had decreased ATP levels, viability and fecundity, as compared to unexposed worms ^5,6^. Thus, as a first approach, we quantified ATP concentration in worms recovered from the intestines of *Chat^LoxP/LoxP^* and *Chat^LoxP/LoxP^;Villin-Cre^ERT2^* mice, as a proxy of their global fitness. No significant difference was found in ATP levels of worms from mice of either genotype recovered at 20 days post infection. In contrast, by day 40, significantly increased ATP levels were found in worms from mice with ACh-deficient tuft cells as compared to control *ChAT^LoxP/^*^LoxP^ littermates, suggesting decreased fitness of worms obtained from an environment containing ACh (Figure 5B).

To confirm this finding, we assessed worm fecundity *ex vivo*, in the presence of controlled concentrations of ACh and levamisole, one of the most common cholinergic anti-helminth drugs ^18^. We quantified egg production by defined numbers of female *H. polygyrus* adult worms, freshly recovered from wildtype infected mice 14 days post-infection to ensure that the worms are present in the gut lumen and not yet altered by luminal ACh. When ACh was added to the culture medium, egg production was significantly decreased, as compared to untreated worms, and the degree of inhibition by ACh was comparable to that obtained after treatment with identical concentrations of levamisole (Figure 5C). These data indicate that worm exposure to ACh causes decreased viability and fecundity, with an efficacy similar to that of a well-established anti-helminth drug.

### Luminal ACh targets worm physiology via their muscarinic acetylcholine receptors

To identify the mechanisms underlying the effect of ACh on worm physiology, we assessed worm fecundity ^5,6^ in the presence of ACh combined with different inhibitors of cholinergic signalling. As above, the addition of ACh to the medium resulted in reduced numbers of faecal eggs. To investigate whether worm AChE may counteract the effects of luminal ACh produced by the host, we combined ACh with neostigmine, an AChE inhibitor. Interestingly, the presence of neostigmine significantly potentiated the inhibitory effect of ACh on egg production by the worms (Figure 5D), strongly suggesting that worm-secreted AChE facilitates helminth survival in their host by degrading luminal ACh produced by host tuft cells.

When ACh was combined with either atropine or mecamylamine, which inhibit muscarinic and nicotinic ACh receptors, respectively, we observed a strong reduction of the effect of ACh on egg production in the presence of atropine while it was essentially intact in the presence of mecamylamine (Figure 5D). This implies that the effects of ACh are mediated by muscarinic ACh receptors expressed by the worms. Finally, none of the drugs had significant effects on egg production when used alone (Figure 5D). Altogether, these data argue that, in addition to their sentinel function in initiating type 2 immune responses, intestinal tuft cells act as effectors of such responses by releasing into the host lumen non-neuronal acetylcholine that directly interferes with helminth physiology through their muscarinic AChRs (Figure 6).

**Figure 6:**
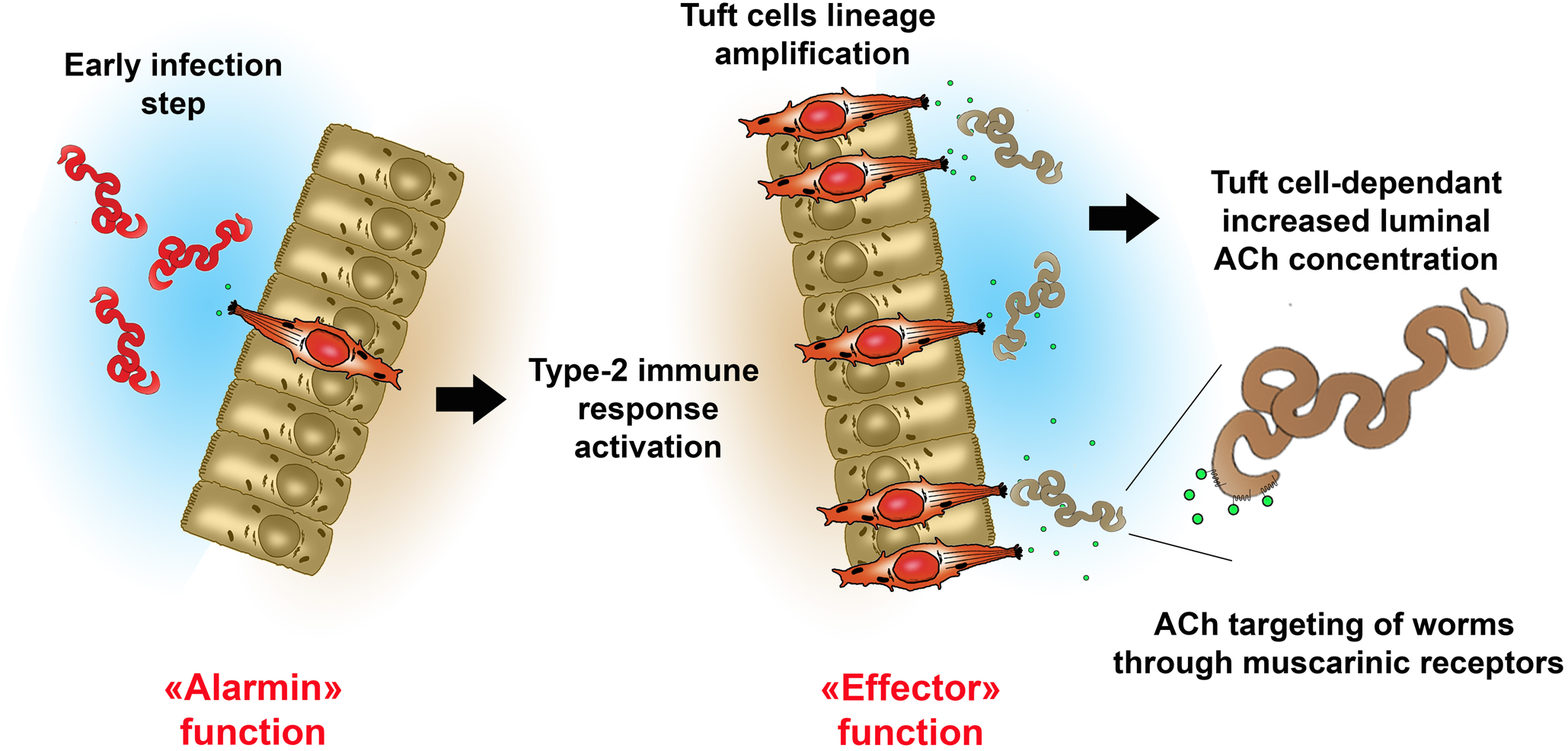
Scheme illustrating the new tuft cell effector function during helminth parasite infection. After initial detection of helminths, tuft cells (red) initiate a type-2 immune response through alarmin secretion, as previously described, eventually leading to mucosal remodelling and drastic tuft cell amplification. Newly produced tuft cells are able to release ACh (green dots) into the lumen to target helminths in a helminth muscarinic receptor-dependant way.

## DISCUSSION

We report here a novel effector function for intestinal tuft cells that justifies, at least in part, the dramatic amplification of this cell lineage in the process of ongoing type 2 responses. This finding may provide the missing link to explain why IL-25 deficiency, abrogating the sentinel function of tuft cells ^8^, causes a less severe phenotype than the complete absence of tuft cells in Pou2f3-deficient mice ^1^.

We also identified slight but significant consequences of the tuft cell ACh deficiency on the numbers of Gata3^+^ immune cells in the *lamina propria* and tuft cells in the intestinal epithelium of naive mice. This suggests additional roles of tuft cell-derived acetylcholine in fine tuning ILC2 homeostasis. Indeed, expression of acetylcholine receptors (AChR) has already been reported on ILC2s, and ACh administration stimulates ILC2 responses after *N. brasiliensis* infection, with higher numbers of total ILC2s, increased percentages of IL-5 producing ILC2s and increased production of type 2 cytokines ^24^, suggesting the possibility that tuft cell-derived ACh might contribute to the regulation of ILC2s. However, since a strong type 2 immune response was established during infections in the absence of tuft cell-derived ACh, we did not investigate this in more detail.

Instead, we focused our study on the role of tuft cell-derived ACh during type 2 immune responses. We report tuft cell-restricted expression of the ChAT enzyme for biosynthesis of ACh, as well as the presence of ACh specifically in tuft cells as compared to other intestinal epithelial cells. Moreover, ACh is present in the gut lumen, with a dramatic increase during type 2 immune responses. Finally, we demonstrate a direct effect of ACh on helminth physiology through their muscarinic AChRs, which is counteracted by parasite-secreted AChE enzymes.

Thus far, the roles played by tuft cell-produced ACh had been mostly studied in the airways. In the mouse tracheal epithelium, tuft cells are capable of sensing bitter compounds present in the airway lining fluid and use cholinergic signalling to neighbouring nerve endings to cause an aversive reflex consisting of a reduced breathing frequency ^10^. Tuft cells also participate in the regulation of bacterial populations present in the airway. These cells can be activated by bacterial quorum sensing molecules (QSM) - used by microbes to evaluate their own population density - such as 3-OxoC_12_-HLS and use ACh signalling to cause respiratory changes ^16^. In addition to respiratory reflexes, mucociliary clearance is an important innate protective process to eliminate inhaled pathogens from the airway. Paracrine cholinergic signalling from tracheal tuft cells, activated by bitter compounds, QSM, or other bacteria-derived products, was found to increase mucociliary clearance by airway ciliated cells, as assessed by the transport speed of particles on the tracheal surface, thereby directly linking chemosensation of bacterial signals with innate defence ^14,17^. Previous studies with reporter mice also reported expression of ChAT ^11^ and of the Tas2r143 bitter taste receptor ^25^ in a subset of tuft cells from the urogenital tract. Furthermore, intraurethral application of a bitter compound regulated bladder activity in rats ^11^.

Previous studies using Chat-GFP reporter mice reported heterogeneity in GFP expression in trachea and urethra Villin-immunoreactive tuft cells ^10,11^. Interestingly, we found co-expression of the endogenous *Chat* mRNA and Dclk1 protein in nearly all analysed cells, suggesting either different regulation of ChAT expression in the tuft cells from distinct organs, or incomplete detection of ChAT expression in genetically engineered reporter mice.

In contrast with these previous studies in the airway and urogenital tracts, we report here a cholinergic function of small intestinal tuft cells with a direct effect of tuft cell-derived acetylcholine on helminth parasites. However, this may not be the unique mode of action of tuft cell acetylcholine in the small intestine and additional cholinergic mechanisms, for instance paracrine signalling to other epithelial cells or nerve cells, as reported in the airway ^10,14^, may also occur. Moreover, additional ACh sources, such as ILC2s, exist in the gut mucosa, and contribute to ILC2 numbers inflation and anti-helminth immunity ^26^.

Luminal ACh production by tuft cells in the context of a type 2 immune response illustrates how different epithelial subsets constitute integral parts of this process. Signalling through epithelial IL-4Rα receptors causes specific amplifications of two epithelial cell lineages. On the one hand, goblet cells not only contribute to the so-called weep-and-sweep response by increased mucus production; they also express *de novo* the Retnlβ peptide that directly interacts with the helminth’s ability to feed on host tissues, and thus decreases their fitness and fecundity ^4–6^. On the other hand, the tuft cell lineage also undergoes dramatic expansion and releases ACh into the gut lumen that also interferes with worm physiology, resulting in reduced global viability and fecundity.

The property of some parasitic helminths to secrete AChE isoforms has long been identified ^27,28^ and, as this is specific to parasitic species, as opposed to free living helminths ^29^, it was thought to confers a selective advantage to the worms in the crosstalk with their host. Thus far, hypotheses on the functions of helminth secreted AChEs have mostly focused on the inhibition of host-intrinsic cholinergic signalling, involved in smooth muscle contraction, mucus production by goblet cells, fluid secretion by enterocytes or cholinergic regulation of the immune system behaviour ^29,30^, although colocalization of AChEs secreted by helminths in the gut lumen and host ACh signalling to muscle or immune cells has not been demonstrated. Since we now demonstrate that, (i) in the context of a type 2 immune response, intestinal tuft cells produce ACh into the gut lumen, (ii) ACh has direct effects on *H. polygyrus* worms, reducing their global fitness and fecundity, (iii) this effect is potentiated upon inhibition of worm-derived AChEs, and (iv) mice with ACh-deficient tuft cells have increased worm burden and faecal eggs, it is tempting to speculate that helminth-secreted AChE into the gut lumen primarily serves to degrade luminal acetylcholine produced by host tuft cells.

As parasitic helminths co-evolved with their hosts, it is plausible that the use of different defence mechanisms, relying on distinct host cell types, such as ACh and Retnlβ, produced by tuft and goblet cells, respectively, renders the task more challenging for parasites to counteract host immunity. In this view, it might be interesting to combine both ACh and Retnlβ host deficiencies to investigate possible synergistic action of these two molecules on different helminth parasite species.

Our study on the cholinergic function of intestinal tuft cells thus reveals a novel function for these cells. Hence, we propose that tuft cells have two distinct but complementary functions in the context of the host defence against helminth infections. In the naïve mucosa, tuft cells are a rare epithelial subset functioning as a sentinel to initiate type 2 immune responses upon parasitic infections. The new finding we now document is that the amplification of the tuft cell lineage, occurring in the context of an ongoing type 2 immune response, underlies a second function as a cholinergic effector cell that directly contributes to parasitic helminth clearance by compromising their global fitness.

Future studies will be needed to evaluate the therapeutic impact of these findings in refining the common anti-helminth treatments of humans or livestock, such as cholinergic agonists or inhibitors of AChEs, with a focus on a luminal mode of action of these drugs.

## Acknowledgments

We acknowledge Prof. M. Selkirk for scientific discussions, Dr. E. Valjent for technical inputs and reagents, Dr. I. Matsumoto for the Pou2f3-deficient mouse line, the RAM-iExplore, RAM-PCEA animal facilities and Luc Forichon for maintenance of mouse colonies, and the Montpellier RIO Imaging (MRI) facilities. This work was supported by a Wellcome Trust Collaborative Award (Ref211814) to PJ, CB, ED, TM and RM, Agence Nationale de la Recherche (ANR-17-CE15-0016-01 and ANR-21-CE15-0017-01 to P.J.), Institut National du Cancer (INCA_2018-158 to P.J.), the PJ team is “Equipe Labellisée Ligue contre le Cancer”; M.N. was supported by the Labex EpiGenMed (an “Investissements d’avenir” program ANR-10-LABX-12-01) and the Wellcome Trust Collaborative Award (Ref211814).

## Author contributions

Conceptualization, P.J., F.G. and R.M.; Methodology, M.D., F.H., C.J., A.T. and S.T.; Investigation, M.D., F.G., F.H., I.G., C.J., S.H. E.T., A.G., and C.C.; Writing – Original Draft, P.J.; Writing – Review & Editing, F.G., R. M. M., C.B., E.D., T.N.M.; Funding Acquisition, R.M.M., P.J., C.B., E.D. and T.N.M.; Resources, S.B. and C.C.; Supervision, P.J., F.G., R.M.

The authors declare no competing interests.

## Material availability

All unique/stable reagents generated in this study are available from the lead contact with a completed materials transfer agreement.

## Figure titles and legends

**Supplementary data Figure 1: Expression of the *Chat* gene in tuft cells throughout the mouse intestine**. Representative *in situ* hybridisation for *Chat* mRNA (red)-coupled with immunofluorescence detection of Dclk1 (green) in Dclk1-expressing tuft cells according to their localisation on the intestinal tract. Scale bar = 10 µm for all panels. Arrows point at tuft cells positive for the Chat signal. Each line shows overlay signals (left), and individual signals in grey levels.

**Supplementary data Figure 2: Efficient Cre^ERT2^ expression in all intestinal epithelial cells, including tuft cells, in the *Chat*^LoxP/LoxP^*;Villin-Cre^ERT2^* mouse strain, after tamoxifen treatment.** Immunostaining for Dclk1 (green) and the ER domain of the chimeric Cre^ERT2^ protein (red), showing nuclear translocation of the Cre^ERT2^ protein in all epithelial cells. Arrows point at tuft cell nuclei. Scale bar = 10 µm.

**Supplementary data Figure 3: Intestinal epithelial *Chat* deficiency does not cause drastic changes in intestinal epithelium cellular composition.** Goblet, Paneth and enteroendocrine cells were counted in naive mice, 10 days after recombination of the *chat* locus. Alcian blue staining was used to highlight goblet cells, and IHC against lysozyme and Insm1, to reveal Paneth and enteroendocrine cells, respectively. Data represent means ± SD of n = 50 villus – crypt axes of each biological replicate (n ranging from 5 to 6; after a Mann Whitney test).

**Supplementary data Figure 4: Expression levels of *Pou2f3* and *Gata3* mRNA in wild type and *Chat*-deficient mice in naive and *H. polygyrus*-infected states.** qRT-PCR experiments were performed either in *Chat^LoxP/LoxP^* control (white bar) or *Chat^LoxP/LoxP^;VillinCre^ERT2^* tamoxifen-treated mice (grey bar), in naive, and *H. polygyrus*-infected mice, 20 and 40 days post-infection. Data represent means ± SD of the different biological replicates (n ranging from 7 to 8 biological replicates, * = p < 0.05, ** = p < 0.01, *** = p < 0.001 after a Mann Whitney test).

**Supplementary data Figure 5: IL-13 stimulation does not cause elevated ACh concentrations in intestinal organoids culture medium**. Relative ACh quantification in the culture medium of either untreated (white bar) or IL13-stimulated (grey bar) intestinal organoids derived from a C57BL/6 mouse after five days. Data represent means ± SD of the different biological replicates (n ranging from 6 to 8 biological replicates, after a Mann Whitney test).

## Materials and Methods

### Ethical statement

All the mice were bred and maintained in an SOPF animal facility. All animal experiments were conducted in accordance with the French Ministry for Education and Research, regarding the care and use of animals for experimental procedures, under the 2022020111554986, and 2019031411197134 Apafis references. Mice were analysed between 8 and 12 weeks of age, regardless of the sex. Cohorts of controls and deficient mice were obtained from littermates. No statistical method was used to predetermine sample size and the experiments were not randomized. Unless otherwise stated, the investigators were not blinded to allocation during experiments and outcome assessment.

### Animal strains and procedures

The conditional *Chat^LoxP/LoxP^* allele,(Chattm1Mlt, ^22^) was kindly provided by Sylvie Berrard and crossed with the intestinal epithelium-specific, tamoxifen-inducible, *Villin-Cre^ERT2^* mouse strain (Tg(Vil1-cre/ERT2)23Syr, ^23^). The Pou2f3-deficient mouse strain was provided by H. Matsumoto (Pou2f3tm1Abek, ^31^). Cre-mediated recombination was achieved with a daily intraperitoneal injection of 1 mg of tamoxifen (Sigma), for 5 consecutive days. For *H. polygyrus* infection experiments, mice were inoculated orally with 200 infective L3, as previously described ^5^. Infection parameters were monitored according to the number of faecal eggs, or number of intestinal adult worms from day 10 to day 40 post infection. Samples of luminal intestinal contents were obtained from naïve or infected mice, according to the following surgery procedure. After 4 hours of fasting, mice received subcutaneously a single dose of buprenorphine (0.1 mg/kg body weight), and were anesthetized 30 min later with isoflurane (3 to 2%). After local lidocaine treatment, the abdominal cavity was opened, and an intestinal loop was ligatured. This intestinal loop was filled with up to 400 µl of a PBS solution. The loop content was recovered after 30 min, after which mice were euthanized.

### Collection of adults *H. polygyrus* and in vitro culture for treatment with cholinergic drugs

Adult *H. polygyrus* worms were collected from the intestinal lumen of infected mice at 14 dpi to evaluate the effect of cholinergic drugs on eggs release and worm fitness. Worms were washed with PBS containing penicillin (5U/ml)/streptomycin (5μg/ml) /gentamicin (1%), counted, and 20 to 25 female worms per well were incubated at 37°C, 5% CO2, in 200µL RPMI containing antibiotics. Adult *H. polygyrus* worms were then treated with the following drugs: Acetylcholine chloride (Sigma, A6625) and Levamisole hydrochloride (Sigma, L0380000) at 10µM or 100µM, Neostigmine Bromide (Sigma, N2001), Atropine (Sigma, A0132), and Mecamylamine hydrochloride (Sigma, M9020) at 5µM. After 24h, culture media were mixed with saturated NaCl solution (in a 50/50 volume ratio) to count eggs. Worms were collected and used as substrate for ATP level quantification, using the ATPLite kit (6016943; Perkin Elmer), according to the manufacturer’s instructions.

### Cell sorting experiments

Single intestinal epithelial cells were obtained from small intestines after incubation in ice cold 30mM EDTA (Sigma) in HBSS pH 7.4 (Life Technologies) for 20 min. Tissues were then vigorously shaken in DMEM (Life Technologies) supplemented with 10% FBS (Sigma), with 100μl of Dispase (BD Biosciences), and 100μl of DNase I at 2,000 Kunitz (Sigma). After filtration on a 40 µm mesh, single cell suspensions were incubated with phycoerythrin rat anti-mouse Siglec-F antibody (BD Pharmigen, 552126), and FITC rat anti mouse EpCam antibody (17-5791-82, ebiosciences) for 30min at 4°C, and washed with HBSS and resuspended in appropriate volume of HBSS pH 7.4 supplemented with 5% FBS before staining with 7-aminoactinomycin D (Life Technologies) to exclude dead cells. Siglec-F+ live cells were sorted using a FACSAria (Becton Dickinson), directly in RLT lysis buffer (Qiagen) for subsequent RNA extraction, or methanol for ACh quantification assay.

### Acetylcholine quantification

A volume of 0.2 mL of intestinal lavage fraction were supplemented with 0.8 mL pure methanol. The sample was kept at −80C for 48h to allow protein precipitation. Next, 2-Morpholinoethansulfonic acid (Cat. # 341-01622; Dojindo, Tokyo, Japan) was added as quality control standard at 0.5 μM final concentration (FC) and the samples were then supplemented with formic acid to 1% FC. The samples were vortexed vigorously for 1 min and then centrifuged for 10 min at 20.000xg to precipitate proteins. A volume of 0.9 mL supernatant was then loaded on the Captiva EMR plate (Agilent, Cat.# 5190-1001) assembled on the Vacuum Manifold (Agilent, Cat.# A796) together with the Deep Well collection plate (Agilent, Cat.# A696001000).The flow-through fractions were first dried using a Speedvac concentrator and then resuspended in 200µL of water. Calibration curve was established by diluting defined amounts of acetylcholine in the same matrix as the samples (generated by intestinal lavage of KO mice). Protein quantities from each sample were measured using the BCA kit (Thermo Scientific) according to manufacturer’s recommendations. Briefly, protein pellets were solubilized in 0.3 mL of 2% sodium dodecylsulfate solution in water. Two uL of sample was then used to estimate the protein content of each sample.

One microliter of resuspended Captiva flow-through fraction or 1 uL of standard were injected on LC-MS systems consisting of UHPLC (Agilent, 1200 Infinity II Biocompatible) coupled to triple quadrupole MS (Agilent, 6495C). The samples were analysed using in MRMs acquisition mode. Following transition were established to quantify acetylcholine: 147.1 → 43.0, 147.1 → 88.1, 147.1 → 87.1. The analytical column was UPLC Discovery™ column HS F5-3 (Cat. # 567503-U; Sigma Aldrich). Mobile phases were composed as follows: A: 99.9% water, 0.1% formic acid (Sigma Aldrich, Cat. # 33015); B: 99.9% acetonitrile (Biosolve BV, Cat.# 001204102BS), 0.1% formic acid. The gradient was: 0 min (100% A), 2 min (100% A), 5 min (75% A), 11 min (65% A), 15 min (5% A), 25 min (5% A), 25.10 min (100% A), 35 min (100% A). Flow rate was 0.25 ml/min and the temperature column was set to 40°C. The following source parameters were used: Gas Temperature: 150°C; Gas Flow: 11 L/min; Nebulizer: 40 psi; Sheath Gas Temperature: 400°C; Sheath Gas Flow: 12 L/min; Capillary Voltage (neg. mode): 4000 V; Capillary Voltage (pos. mode): 4000 V; Nozzle Voltage: 500 V; IFunnel High Pressure RF Pos: 100 V Neg: 50 V; IFunnel Low Pressure RF Pos: 100 V Neg: 50 V.

Following the analysis, peak integration was conducted using Agilent Masshunter Quantitative Analysis software (for QQQ). The absolute quantification of acetylcholine was calculated by applying the peak area from each sample to the acetylcholine calibration curves.

### 3D organoid culture methods

Organotypic cultures from mouse small intestinal epithelial cells were prepared and propagated as previously described ^32^. In brief, a third of the small intestine was recovered, washed with PBS solution supplemented with antibiotics. Tissues were then cut in small pieces and incubated in 2mM EDTA (Sigma-Aldrich) in PBS for 30 minutes on ice. After vigorous pipetting, crypts were isolated by filtration on a 70 μM cell strainer and resuspended in DMEM/F12 supplemented with an antibiotic mix, 10mM Hepes and 2mM L-Glutamine, before to be plated in Matrigel (Corning), in a DMEM culture medium supplemented with: B-27 supplement (Gibco), N-acetylcysteine (1.25mM), EGF (50 ng ml−1) (Peprotech), Noggin (100 ng ml−1) (R&D system) and R-Spondin (500 ng ml−1) (Peprotech). Cultures were replated every five days.

### RNA extraction and PCR

Total RNA from intestinal tissues were isolated using TRIzol (Life Technologies). RNeasy Mini and Micro Kit columns (Qiagen), were also used for RNA purification from organoid cultures or cell-sorted experiments, respectively. Sorted-cell RNA were further amplified using the Arcturus RiboAmp Plus kit (ThermoFischer Scientific, KIT0501). Reverse transcription was performed with 1 μg of purified RNA using Transcriptor First Strand cDNA synthesis KIT (Roche) according to the manufacturer’s instructions. Real-time quantification was performed in triplicate with a LightCycler480 (Roche) using LightCycler 480 SYBR Green I Master (Roche) on 5ng of RT product using the average Ct of Gapdh and Hprt as internal loading controls, and ΔΔCt method was used for calculating relative expression.

### Fluorescent immunohistochemistry or *in-situ* hybridisation on paraffin-embedded tissue

Tissue dissection, fixation, and immunohistochemistry on thin sections of paraffin-embedded tissue were performed essentially as described previously ^33^. Epitope retrieval was achieved by boiling in 10 mM in sodium citrate (pH 6.4) during 20 minutes. Primary antibodies used in this study were incubated ON at 4°C, and are listed according to the STAR method protocol. Slides were then washed twice with 0.1% PBS-Tween (Sigma-Aldrich) before incubation with fluorescent dyes-conjugated secondary antibodies (Jackson ImmunoResearch Laboratories, Inc.) and Hoechst at 2 μg ml−1(Sigma-Aldrich) in TBS–Triton X-100 0.1% (Sigma-Aldrich), or HRP-conjugated secondary antibodies, revealed with DAB (Sigma). Slides were mounted in FluoroMount (Sigma) or Pertex (Histolab), for fluorescent or visible imaging, respectively. For mRNA *in situ* hybridisation, tissues were hybridised with a probe targeting the *Chat* mRNA (Cat No. 408731, ACDBio). Slides were processed according to the manufacturer’s instructions until probe revelation, after which immunofluorescence detection of Dclk1 was performed following the methodology described above.

### Microscopy and imaging

Fluorescent pictures were acquired at room temperature on an AxioImager Z1 microscope (Carl Zeiss, Inc.) equipped with a camera (AxioCam MRm; Carl Zeiss, Inc.), EC Plan Neofluar (5X NA 0.16; 10X NA 0.3; 20X 0.5 NA; 100X NA 1.3) and Plan Apochromat (40X NA 0.95; 63X NA 1.4) lenses, apotome Slider system equipped with an H1 transmission grid (Carl Zeiss, Inc.), and Zen software (Carl Zeiss, Inc.). Post-treatment of pictures (level correction), annotations, and panel composition were performed using the Photoshop (Adobe) or Zen (Carl Zeiss) software. Bright-field immunohistochemistry pictures were taken at room temperature on an Eclipse 80i microscope (Nikon) with Plan Fluor (10X NA 0.3; 20X NA 0.5; 40X NA 0.75; and 60X NA 0.5–1.25) lenses (Nikon) and a digital camera (Q-Imaging Retiga 2000R with a Q-Imaging RGB Slider), with Q-Capture Pro software (Nikon). Stained slides were also imaged with the Nanozoomer device (Hamamatsu), visualised and annotated with the NdpView Software (Hamamatsu).

### Statistical analyses

Prism software was used for descriptive statistical analyses. For infection monitoring, sample (n) was defined as the number of eggs per gram of faeces per mouse or the number of eggs per well. For histological data quantification, sample (n) was defined as the mean of number of cells per crypt– villus unit per mouse. Unless otherwise stated, 50 crypt–villus axes were counted per histological sections from 5 to 10 mice of each genotype or condition. As the data were not normally distributed, a two-tailed Mann–Whitney U-test was used to calculate the P values. Results are shown as histograms representing means as centre values and standard deviation as error bars for each genotype or conditions, where individual data point are plotted.

## STAR Methods

### KEY RESOURCES TABLE

The table highlights the reagents, genetically modified organisms and strains, cell lines, software, instrumentation, and source data **essential** to reproduce results presented in the manuscript. Depending on the nature of the study, this may include standard laboratory materials (i.e., food chow for metabolism studies, support material for catalysis studies), but the table is **not** meant to be a comprehensive list of all materials and resources used (e.g., essential chemicals such as standard solvents, SDS, sucrose, or standard culture media do not need to be listed in the table). **Items in the table must also be reported in the method details section within the context of their use.** To maximize readability, the number of **oligonucleotides and RNA sequences** that may be listed in the table is restricted to no more than 10 each. If there are more than 10 oligonucleotides or RNA sequences to report, please provide this information as a supplementary document and reference the file (e.g., See Table S1 for XX) in the key resources table.

*Please note that ALL references cited in the key resources table must be included in the main references list.* Please report the information as follows:

- **REAGENT or RESOURCE:** Provide the full descriptive name of the item so that it can be identified and linked with its description in the manuscript (e.g., provide version number for software, host source for antibody, strain name). In the experimental models section (applicable only to experimental life science studies), please include all models used in the paper and describe each line/strain as: model organism: name used for strain/line in paper: genotype. (i.e., Mouse: OXTR^fl/fl^: B6.129(SJL)-Oxtr^tm1.1Wsy/J^). In the biological samples section (applicable only to experimental life science studies), please list all samples obtained from commercial sources or biological repositories. Please note that software mentioned in the methods details or data and code availability section needs to also be included in the table. See the sample tables at the end of this document for examples of how to report reagents.
- **SOURCE:** Report the company, manufacturer, or individual that provided the item or where the item can be obtained (e.g., stock center or repository). For materials distributed by Addgene, please cite the article describing the plasmid and include “Addgene” as part of the identifier. If an item is from another lab, please include the name of the principal investigator and a citation if it has been previously published. If the material is being reported for the first time in the current paper, please indicate as “this paper.” For software, please provide the company name if it is commercially available or cite the paper in which it has been initially described.
- **IDENTIFIER:** Include catalog numbers (entered in the column as “Cat#” followed by the number, e.g., Cat#3879S). Where available, please include unique entities such as <u>RRIDs</u>, Model Organism Database numbers, accession numbers, and PDB, CAS, or CCDC IDs. For antibodies, if applicable and available, please also include the lot number or clone identity. For software or data resources, please include the URL where the resource can be downloaded. Please ensure accuracy of the identifiers, as they are essential for generation of hyperlinks to external sources when available. Please see the Elsevier <u>list of data repositories</u> with automated bidirectional linking for details. When listing more than one identifier for the same item, use semicolons to separate them (e.g., Cat#3879S; RRID: AB_2255011). If an identifier is not available, please enter “N/A” in the column.

- ***A NOTE ABOUT RRIDs:*** We highly recommend using RRIDs as the identifier (in particular for antibodies and organisms but also for software tools and databases). For more details on how to obtain or generate an RRID for existing or newly generated resources, please <u>visit the RII</u> or <u>search for RRIDs</u>.

Please use the empty table that follows to organize the information in the sections defined by the subheading, skipping sections not relevant to your study. Please do not add subheadings. To add a row, place the cursor at the end of the row above where you would like to add the row, just outside the right border of the table. Then press the ENTER key to add the row. Please delete empty rows. Each entry must be on a separate row; do not list multiple items in a single table cell. Please see the sample tables at the end of this document for relevant examples in the life and physical sciences of how reagents and instrumentation should be cited.

### TABLE FOR AUTHOR TO COMPLETE

*Please upload the completed table as a separate document. **<u>Please do not add subheadings to the key</u> <u>resources table.</u>** If you wish to make an entry that does not fall into one of the subheadings below, please contact your handling editor. **<u>Any subheadings not relevant to your study can be skipped.</u>** (**NOTE:** References within the KRT should be in numbered style rather than Harvard.)*

#### Key resources table

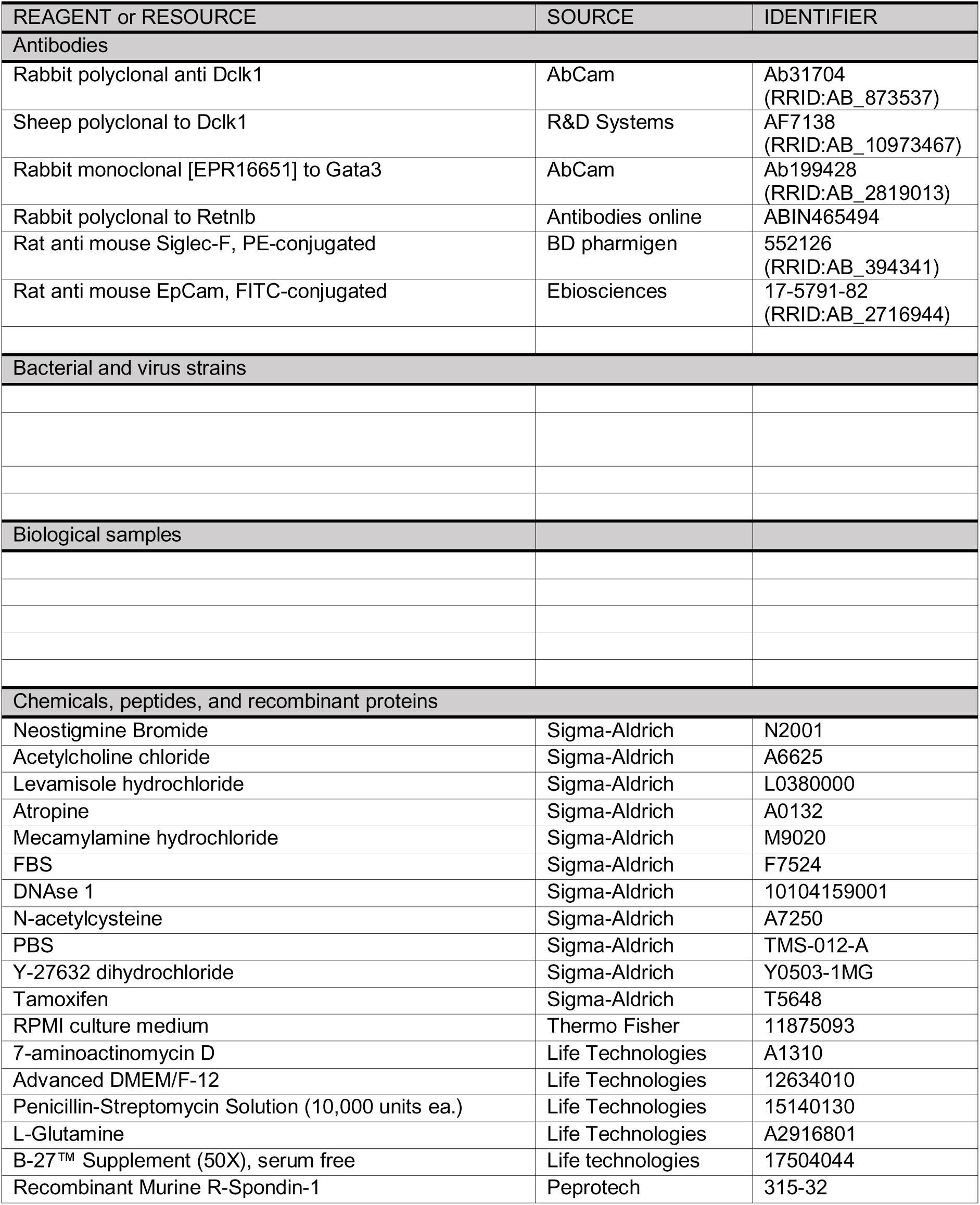
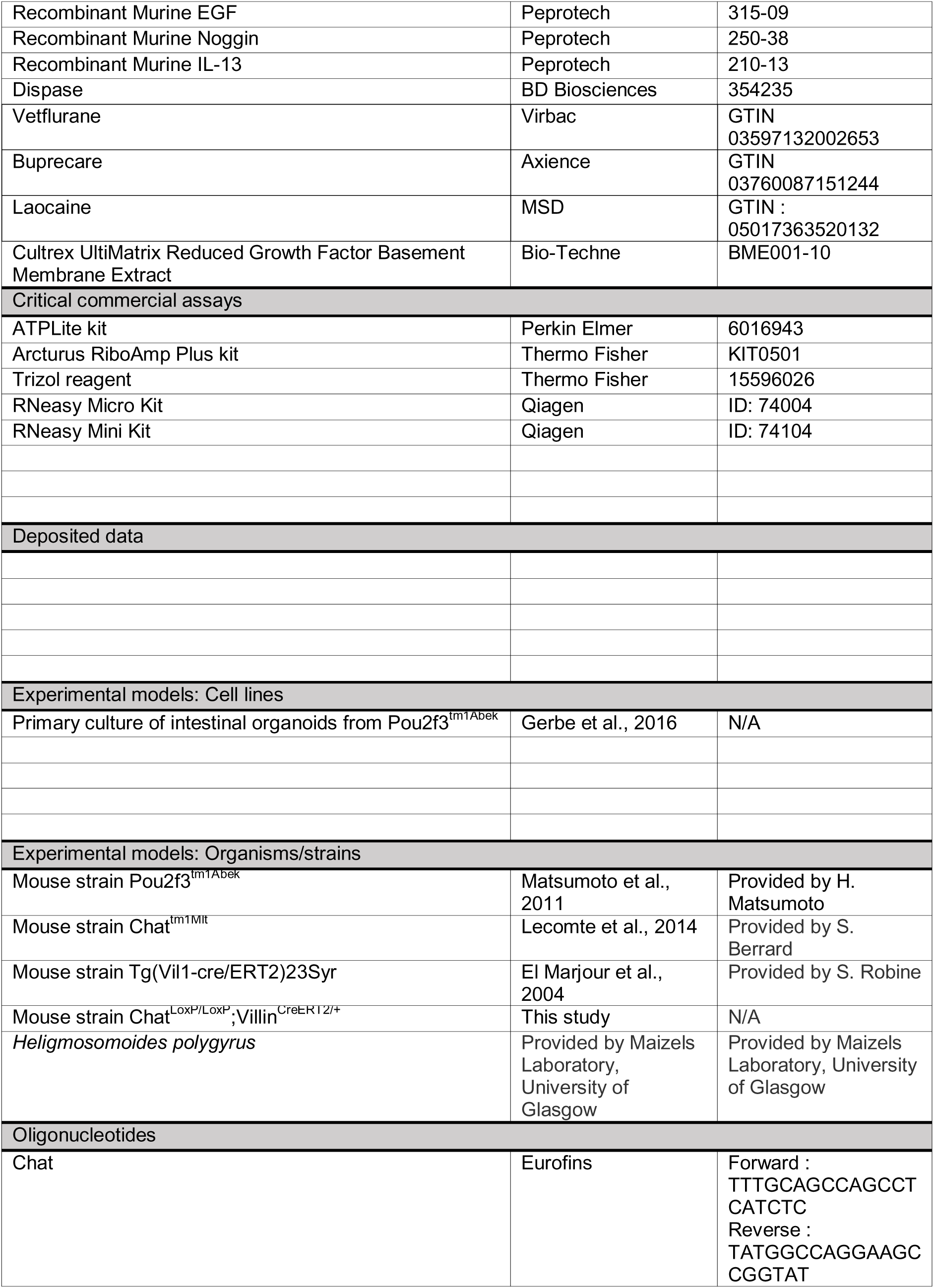
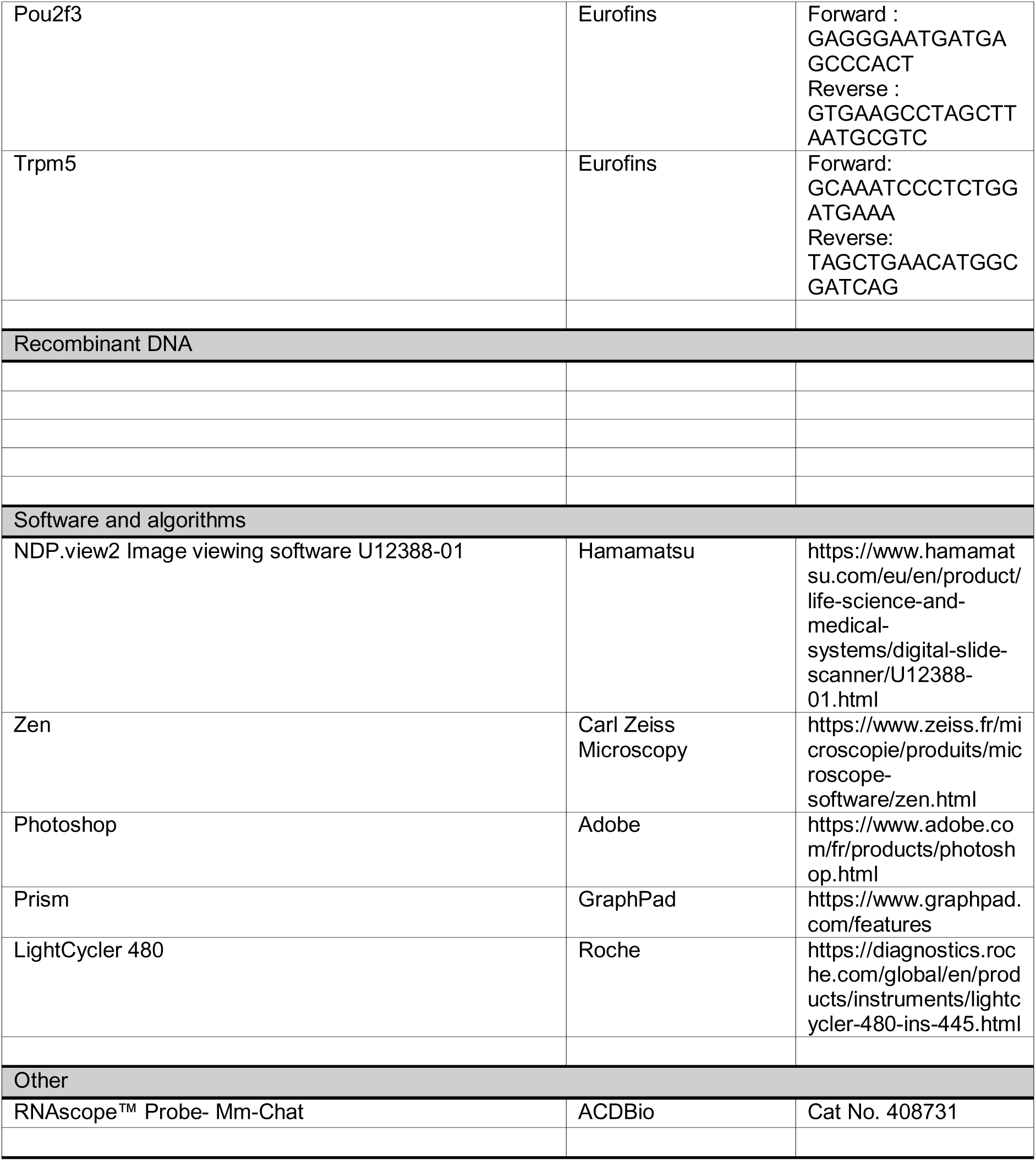

