## Supplementary figures and images for "Tuft cell-derived acetylcholine is an effector of type 2 immunity and directly targets helminth parasites in the gut lumen"

### Supplemental figure 1

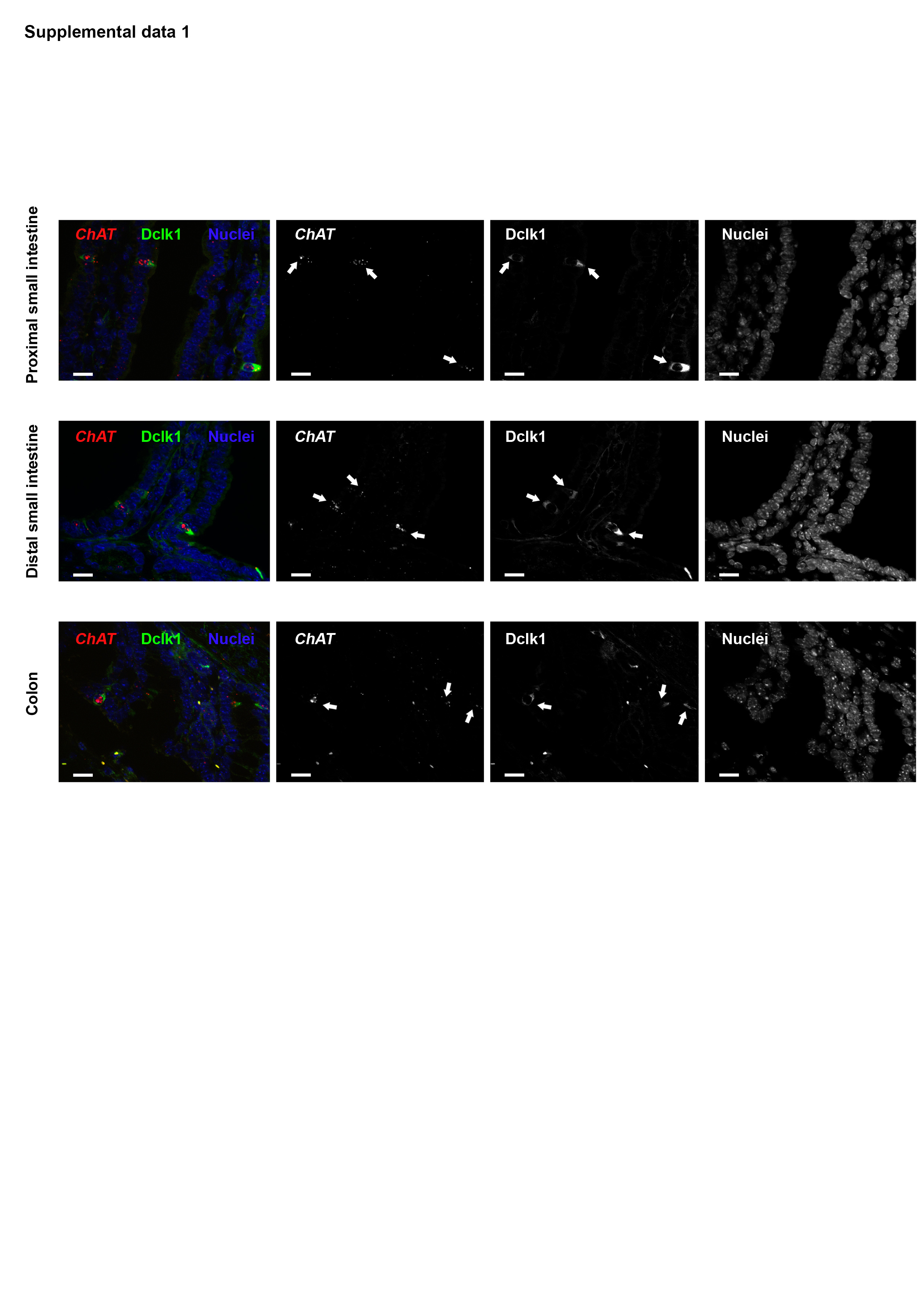

### Supplemental figure 2

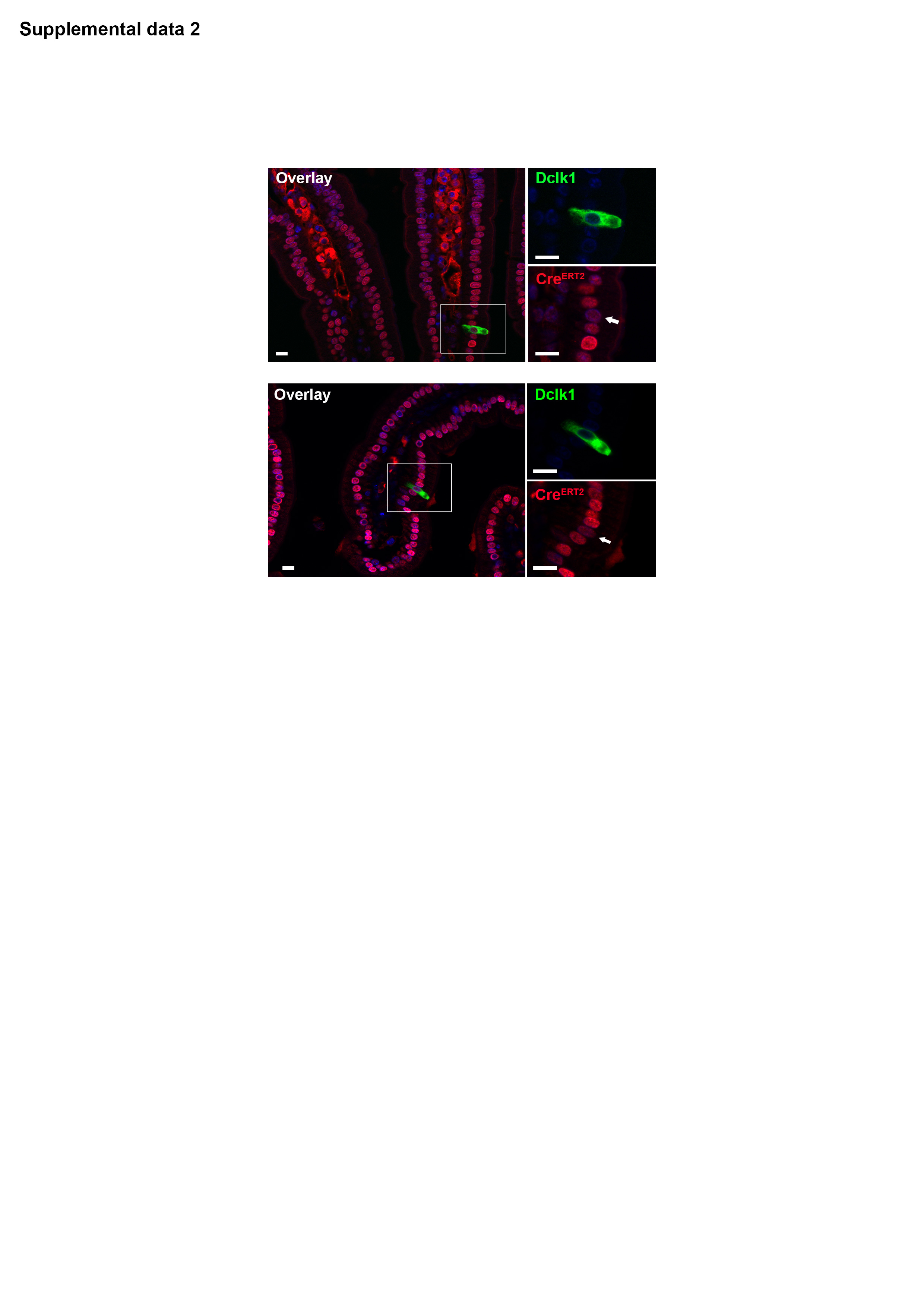

### Supplemental figure 3

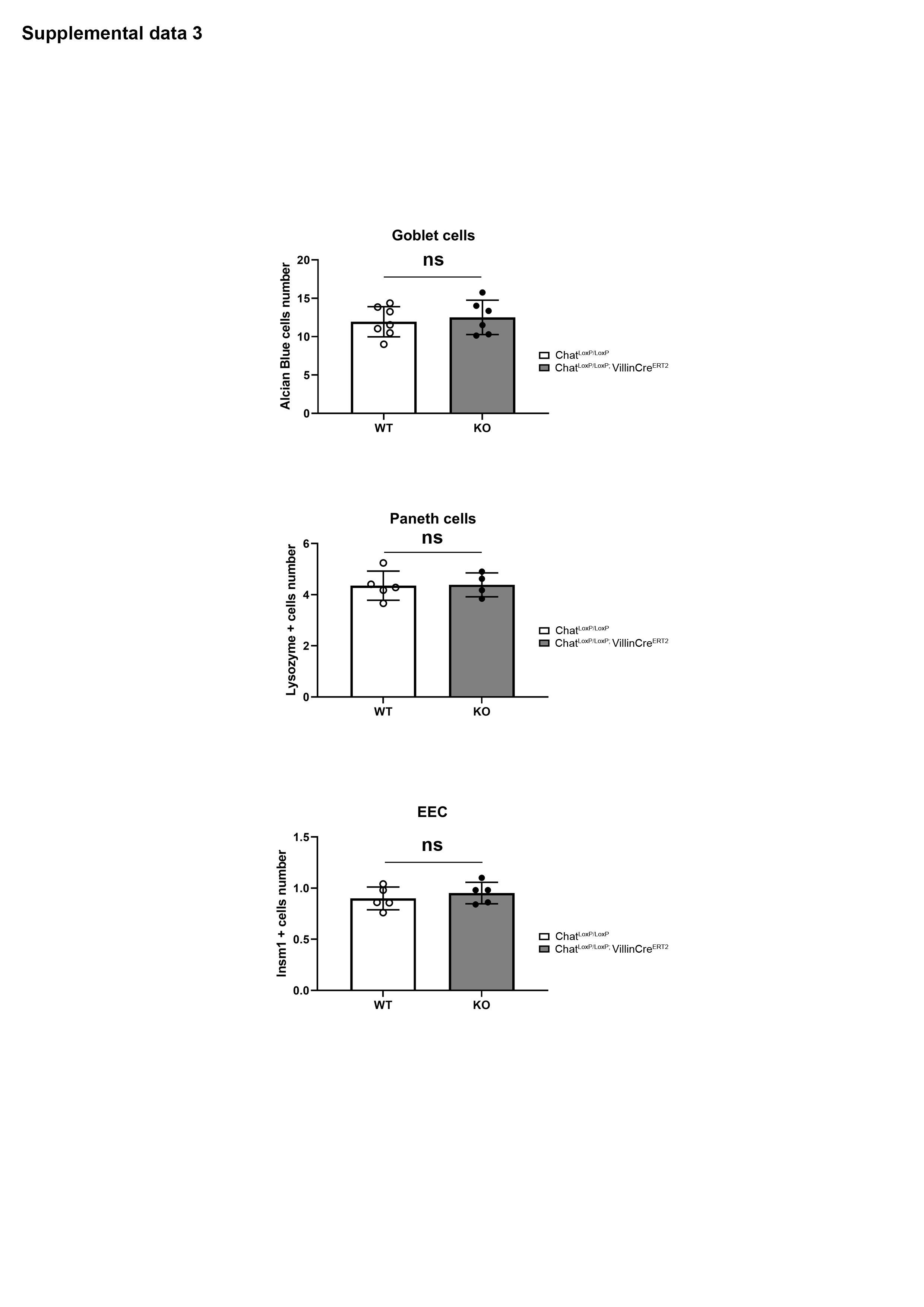

### Supplemental figure 4

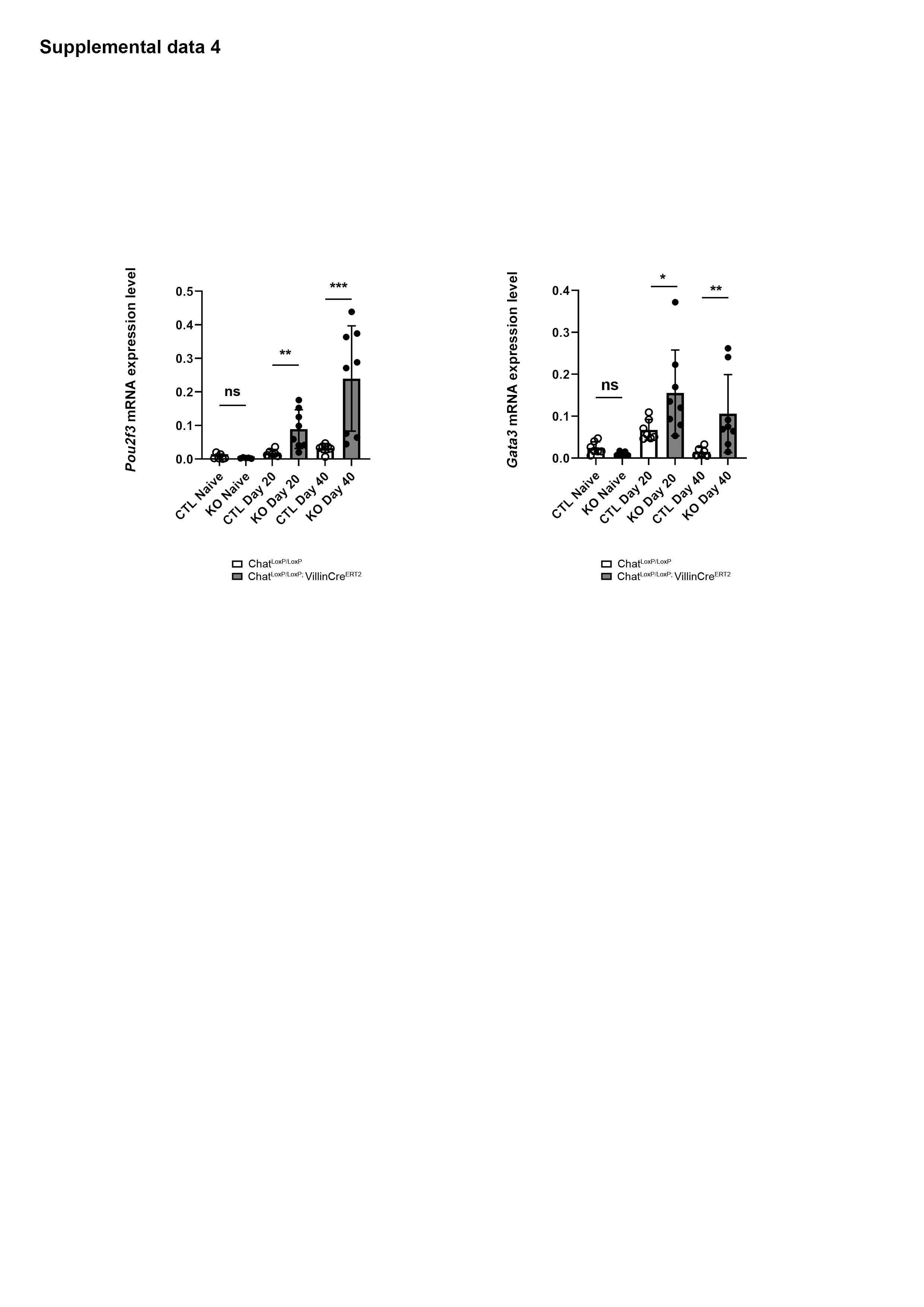

### Supplemental figure 5

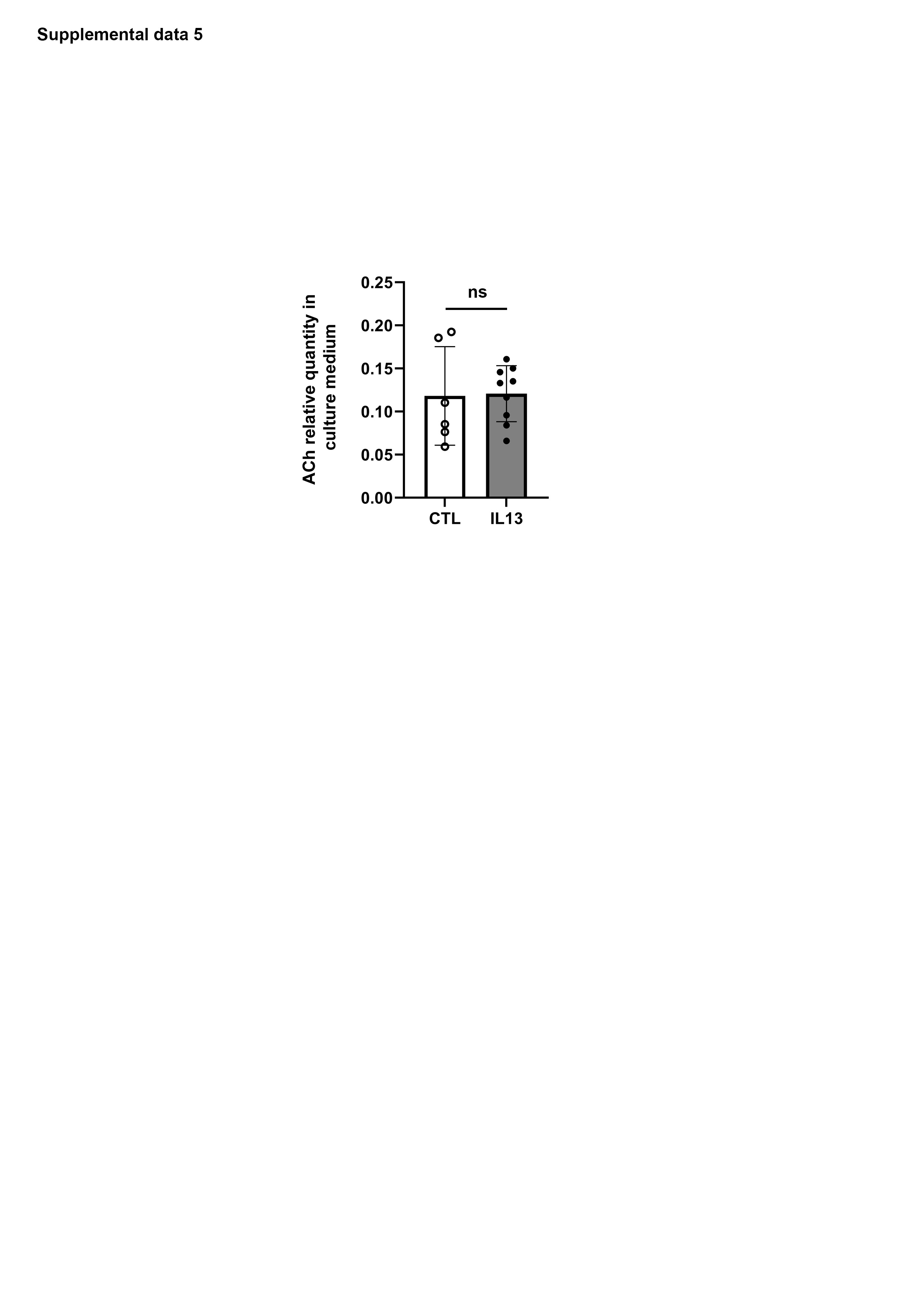
